# Understanding TCR T cell knockout behavior using interpretable machine learning

**DOI:** 10.1101/2024.10.01.616134

**Authors:** Marcus Blennemann, Archit Verma, Stefanie Bachl, Julia Carnevale, Barbara E. Engelhardt

**Affiliations:** Gladstone Institutes, San Francisco, CA 94158, USA; University of California, San Francisco, San Francisco, CA 94143, USA; Biomedical Data Science, Stanford University, Stanford, CA 94305, USA

**Keywords:** Explainable AI, Grad-CAM, machine learning, live cell imaging

## Abstract

Genetic perturbation of T cell receptor (TCR) T cells is a promising method to unlock better TCR T cell performance to create more powerful cancer immunotherapies, but understanding the changes to T cell behavior induced by genetic perturbations remains a challenge. Prior studies have evaluated the effect of different genetic modifications with cytokine production and metabolic activity assays. Live-cell imaging is an inexpensive and robust approach to capture TCR T cell responses to cancer. Most methods to quantify T cell responses in live-cell imaging data use simple approaches to count T cells and cancer cells across time, effectively quantifying how much space in the 2D well each cell type covers, leaving actionable information unexplored. In this study, we characterize changes in TCR T cell’s interactions with cancer cells from live-cell imaging data using explainable artificial intelligence (AI). We train convolutional neural networks to distinguish behaviors in TCR T cell with CRISPR knock outs of CUL5, RASA2, and a safe harbor control knockout. We use explainable AI to identify specific interaction types that define different knock-out conditions. We find that T cell and cancer cell coverage is a strong marker of TCR T cell modification when comparing similar experimental time points, but differences in cell aggregation characterize CUL5KO and RASA2KO behavior across all time points. Our pipeline for discovery in live-cell imaging data can be used for characterizing complex behaviors in arbitrary live-cell imaging datasets, and we describe best practices for this goal.

## 1. Introduction

Since FDA approval in 2017, chimeric antigen receptor (CAR) T cell immunotherapies have proven effective at treating advanced leukemias and lymphomas.^1,2^ CAR T cell therapy uses *ex vivo* modification of native patient T cells to express a chimeric antigen receptor (CAR), capable of binding to surface markers of cancerous cells, to enhance T cell immunological response. T-cell receptor (TCR) therapy is a related method of treatment that uses naturally existing TCRs, protein complexes that bind to a cell’s major histocompatibility complex (MHC), as an alternative to CAR proteins. TCR therapy targets various cancers by recognizing a specific antigen presented by a human leukocyte antigen (HLA) on cancer cell surfaces. This reduces the risk of toxicity associated with CAR T cell therapy, which currently struggles to distinguish between solid cancer cells and normal tissues.^1^ TCR therapy, in contrast, has demonstrated effective responses against multiple solid cancer types such as melanoma and lung carcinoma with reduced off-target effects.^3^

Genetic editing of CAR and TCR T cells with CRISPR-based tools is an emerging approach to engineer improved T cell therapies. CRISPR knock out of the *RASA2* (RASA2KO) or *CUL5* (CUL5KO) genes, for example, has been demonstrated to improve T cell performance against cancer cells *in vitro*.^4^ *RASA2* is a signalling checkpoint in human T cells and increases in response to chronic antigen exposure. TCR and CAR T cells without *RASA2* show better activation, higher cytokine production, and increased metabolic activity, en route to improved cancer cell removal. These RASA2KO T cells also have a survival advantage in mouse models of leukemia and other cancers.^4^ *CUL5* is known to be a negative regulator of the signaling pathways in cytotoxic T lymphocytes. Knocking out *CUL5* has been shown to effectively inhibit tumor growth in mouse studies.^5^ Although these genes have been identified as effective modifications in TCR T cells in *in vivo* mouse studies, understanding the biological mechanisms underlying these positive outcomes remains a challenge due to the complex, multi-scale nature of T cell and cancer cell interactions in humans.

Live-cell imaging is a common approach for evaluating the success of different types of modified T cells. Live-cell imaging with high-resolution 2D imaging from one or more channels, usually a bright field along with fluorescent marker channels, across days at fixed time intervals (e.g., every four minutes) captures the dynamics of co-cultures of cancer and modified T cells. Traditional analyses quantify the total amount of cancer cell-specific fluorescent markers as a proxy of tumor response to treatment.^4–6^ Live-cell imaging has been used to identify dynamic behavior such as morphological changes during T cell killing, or differences in response to liquid or solid tumors, using deep learning methods to segment sequential images.^7^ Even with existing approaches, many questions about dynamic cellular behaviors are difficult to answer.

Computer vision, a subfield of AI, is advancing rapidly in biomedical imaging. Deep learning models, especially convolutional neural networks (CNNs), enable extraction of complex phenotypes from live-cell imaging data. This includes cell segmentation, single-cell tracking, spatiotemporal pattern recognition, and predictive modeling, all of which may be used to study the therapeutic behavior of these modified T cells. Efforts are underway to integrate CNN-driven platforms with patient-derived organoids (PDOs) for personalized drug research, exemplified by projects like OrganoID^8^ and OrBITS.^9^ While these tools are powerful, their prediction processes are black-box and challenging to understand. Interpreting a CNN’s decision-making process should provide important information for researchers attempting to gain biological insights from their live-cell experiments. Explainable AI techniques have emerged that allow researchers to interrogate the features of images that most directly explain deep learning models’ predictions and performance.^10,11^

In this work, we demonstrate the ability of explainable AI to characterize modified T cell behavioral changes under genetic perturbation. We identify phenotypic differences between TCR T cells with beneficial *RASA2* or *CUL5* knock-outs from live-cell imaging data versus TCR T cell negative controls. We use a suite of CNN classifiers trained to predict one of three genetic perturbations captured in live-cell imaging of TCR T cells co-cultured with cancer cells. We use Grad-CAM, an image explainable AI technique that estimates the change in prediction as a function of changes in pixel space, to identify the specific regions in held-out live-cell images that inform prediction for control Safe Harbor KO, RASA2KO, and CUL5KO TCR T cells. Grad-CAM highlights the regions of the image that contribute to classification as each output class. By highlighting regions that contribute to classification decisions, the Grad-CAM interpretation of images allows us to identify the cell-level phenotypic changes associated with each TCR T cell experiment, and we use these interpretable image markers to characterize the distinct T cell behaviors in the three experimental conditions. Our work develops an interpretable deep learning workflow for the analysis of live-cell imaging data, and we show the benefits of our approach by characterizing the differential behavior of SHKO (control), RASA2KO, and CUL5KO TCR T cells.

## 2. Methods

### 2.1. Data Generation

#### 2.1.1. Isolation of primary T cells from healthy donors

Leukopaks from deidentified healthy donors with approved IRBs were purchased from StemCell Technologies. Primary human T cells were isolated with the EasySep Human T Cell Isolation Kit (StemCell Technologies) according to the manufacturer’s protocol. T cells were seeded at a density of 1 million cells per mL maintained in X-Vivo-15 medium supplemented with 5% fetal bovine serum, 50 *µ*M beta-mercaptoethanol, and 10 mM N-acetyl-L-cysteine plus 100 IU/mL of IL-2 and activated with Dynabeads Human T-Activator CD3/CD28 (Gibco) at a 1:1 bead-to-cell ratio.

#### 2.1.2. CRISPR KO in primary human T cells using Cas9–RNP electroporation

T cell transduction was accomplished by adding concentrated lentivirus directly to the T cells 24 hours after activation with Dynabeads Human T-Activator CD3/CD28, 40 *µ*L virus per 1 *×* 10^6^ T cells in X-Vivo-15. At 48 h post-activation, Cas9–sgRNA–RNP electroporation was conducted with the Amaxa P3 Primary Cell 96-well 4D-Nucleofector Kit (Lonza). The safe harbor T cells were targeted using the *AAVS1* sequence GGGCCACTAGGGACAGGAT, the *RASA2* -ablation T cells with the sequence AGATATCACACATTACAGTG, and the *CUL5* -ablation T cells with the sequence ATTGGAGTAAGAGAATCCTA. crRNAs and tracrRNAs were then complexed 1:1 by volume and incubated for 30 minutes at 37C to form sgRNAs. The sgRNAs were then mixed with Cas9 (stock concentration of 40 *µ*M, QB3 Macrolab) at 1:1 by volume for 15 minutes at 37C to produce ribonucleoproteins (RNPs) complexes. After counting, T cells were resuspended in P3 buffer at 1 *×* 10^6^ per 20 *µ*l, mixed with 3 *µ*l of RNPs, and added to a 96-well electroporation plate. Electroporation was performed using using the EH115 protocol and recovered by adding 80 *µ*l T cell medium (X-Vivo-15, Lonza) at 37C for 15 min. Cells were transferred to appropriate culture vessels containing X-Vivo-15 medium supplemented with IL-2 containing 100 IU per mL.

#### 2.1.3. Repetitive stimulation assay

Tumor cells were maintained in a complete RMPI (Gibco) consisting of 1% penicillin-streptomycin (Gibco), GlutaMAX supplement (Gibco) and 10% fetal bovine serum (Corning), and then resuspended in T cell medium. T cells were seeded on top of the cancer cells at a 1:1 E:T ratio with IL-2 at 100 IU mL^*−*1^. Subsequent repeated co-cultures were set up every 48 h. For each co-culture, T cells were counted using the Cellaca MX High-throughput Cell Counter (Revity), percentage of TCR+ cells was measured via flow cytometry, and T cells were replated onto fresh tumor cells every 48 hours maintaining a 1:1 E:T ratio.

#### 2.1.4. In vitro cancer killing assay by TCR T cells

Antigen-specific T cells were co-cultured in X-VIVO-15 plus supplements – 100 IU IL-2 per mL and 1X Glucose (Gibco) – with mKate+ A375 cells pre-seeded in a 96-well flat-bottom plates at a 1:1 E:T ratio. Images were captured every 4 minutes over a 24-hour span using the IncuCyte S3 live-cell imaging platform (Essen Bioscience). The mKate+ object counts for each well were recorded over time.

### 2.2. Model architecture, training, and evaluation

A convolution neural network was trained to identify the TCR T cell genetic perturbation – RASA2KO, CUL5KO, or SHKO – from a single 300 by 300 pixel subsection of each image. The network was trained on images from nine of the replicates, three from each condition, and validated on the remaining three held-out wells. The model consists of a ResNet50^12^ block, a fifty layer residual convolutional neural network (CNN), that feeds into a fully connected linear layer to predict the weights for each class. ResNet50 is a CNN designed for image classification tasks. The first stage consists of 64 7 *×* 7 convolutional filters, followed by four stages of residual blocks. These stages contain filters configured as follows: the first has 64, 64, and 256 filters; the second has 128, 128, and 512 filters; the third has 256, 256, and 1,024 filters; and the fourth has 512, 512, and 2,048 filters. The network ends with a fully connected layer with one neuron per possible output class, which is three in our application. The initial convolutional layer and first layer of each stage uses a stride size of 2, while all other layers use a stride size of 1. The weights and biases of the final, fully connected output linear layer were trained to minimize the cross-entropy loss of the predicted probability of each class, a softplus of the linear output layer, to the true data label.

The untrained parameters of the ResNet50 block were initialized as the parameters of ImageNet.^13^ These weights capture high-level features, such as edges and shapes, allowing us to reach accurate classification faster. The last layer of the model was fine tuned on 12,600 unique frames of our training data, evenly split among the three conditions, with two frames per batch. We use the quarter-sectioned 300 *×* 300 pixel images to minimize the effects of downsampling, as ResNet50 takes as input 224 *×* 224 pixel images and downsamples larger inputs. The brightfield phase images were converted from grayscale to RGB to match the required input parameters of Resnet50. The CNN was fine tuned with the Adam optimizer^14^ for forty epochs with a learning rate of 1 *×* 10^*−*3^. The same procedure and architecture was also used to train a CNN classifier on a subset of frames from between 800 and 9996 minutes (frames 200 through 249 out of 350 total) into the experiment, a total of 1800 images, to evaluate the time dependence of the predictions. The model was trained on an NVIDIA A30 GPU using CUDA, PyTorch, and PyTorch lightning.

To obtain “visual explanations” for the classification of each frame, we applied the gradient-weighted class activation mapping (Grad-CAM) technique^10^ to the model for each frame of the validation set. This technique computes the gradients of the target class score with respect to the feature maps of the final convolutional layer of the network. These gradients are pooled across the convolutional filter to provide a spatial-average importance value for different regions of the input image that contribute to the target class score. Grad-CAM returns an “importance” of each pixel to the final prediction that can be superimposed onto the original images and visually inspected to identify relevant image details. Model training and analysis code is available at https://github.com/25marcusb/Understanding-TCR-T-cell-knockout-behavior-using-interpretable-machine-learning.

## 3. Results

### 3.1. Convolutional neural networks can distinguish between different genetic perturbations from a single frame

We first validate the predictive ability of the trained CNN classifier to distinguish the genetic perturbation given a section of a live-cell imaging frame. The CNN was trained to distinguish between three classes -RASA2KO T cells, CUL5KO T cells, and SHKO T cells. RASA2KO and CUL5KO are known to improve T cell anti-cancer activity after repetitive stimulation.^4,5^ The SHKO T cells, with *AAVS1* knockouts as a negative control, should be “exhausted” after repetitive stimulation, leading to less anti-cancer activity.^4^ On held-out validation data, the model assigns more than 50% probability to the correct class on 2,974 out of 4,200 test images evenly balanced across classes, an accuracy of 71%. The CNN model outperforms a traditional support vector machine (SVM) classifier that predicts the perturbation from cell counts from segmentation, which has a test accuracy of only 50%. We find that the model has consistent precision around 70%, e.g., the fraction of true RASA2KO frames out of the set of all frames predicted to be RASA2KO, across the genetic perturbations (Table 1). However, we observe that the ability to recall the SHKO control condition is much worse than the ability to recall the genetic perturbations (Table 1). While the “confusion” with the safe harbor control indicates that many features of T cell and cancer cell dynamics are maintained after perturbation, the relatively low number of incorrect cross predictions between the two genetic knock-outs suggests the model can differentiate the changes from CRISPR perturbation and be used as a tool to interrogate the different behaviors.

**Table 1.** Full prediction model confusion matrix. The rows represent the true labels for the three experiment types; the columns represent the predicted labels. The last row and column of the matrix are the precision and recall, respectively, for each experiment class label.

|  | Predicted SHKO | Predicted CUL5KO | Predicted RASA2KO | Recall |
| --- | --- | --- | --- | --- |
| True SHKO | 387 | 609 | 373 | 28% |
| True CUL5KO | 67 | 1318 | 13 | 94% |
| True RASA2KO | 114 | 7 | 1268 | 91% |
| Precision | 68% | 68% | 76% |  |

We observe a relationship between the collection time of the image and the ability to accurately classify its genetic perturbations. For the control SHKO images, the model tends to classify early time frames as CUL5KO and later images as RASA2KO (Figure 1). To better understand how time affects classification performance, we trained a *limited-time* model with the same architecture, but we restricted the training data to include only images from frames 200 to 249, between 800 and 996 minutes post culture, around the inflection point of RASA2KO and CUL5KO mis-classification. We find that this limited-time model has higher validation accuracy of 89% on held-out data (also in the same time window), and makes relatively few misclassifications (Table 2). CNN-based prediction again outperforms a cell count based SVM classifier, which has an overall test accuracy of 64%. This model does not generalize well to early time frames, but has above 75% accuracy in the 200 minute periods before and after its training data (Figure 2). The inability to generalize well to early time frames is expected given the lack of differentiation between all three conditions in the early parts of the experiment. More generally, this change in predictive ability over time reveals that genetic perturbations may affect the dynamics and timing of immune and cancer cell interactions.

**Table 2.** Limited time (frames 200 -250) test confusion matrix. Rows represent true labels for the three experiment types; columns represent predicted labels. The last row and column of the matrix are the precision and recall, respectively, for each experiment class label.

|  | Predicted SHKO | Predicted CUL5KO | Predicted RASA2KO | Recall |
| --- | --- | --- | --- | --- |
| True SHKO | 184 | 9 | 6 | 92% |
| True CUL5KO | 1 | 198 | 1 | 99% |
| True RAS2KO | 41 | 1 | 154 | 77% |
| Precision | 81% | 95% | 86 % |  |

**Fig 1.**
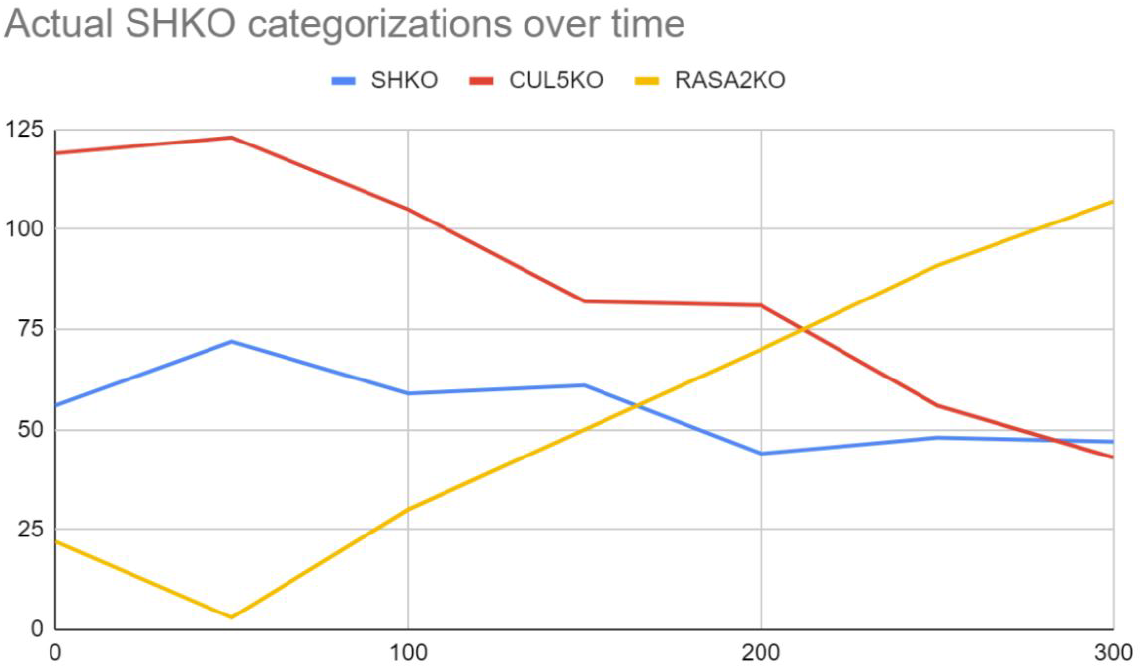
SHKO categorizations over time. Each point corresponds to the 50 frame time bucket starting at that frame (time). A total of 200 images per time bucket are categorized.

**Fig 2.**
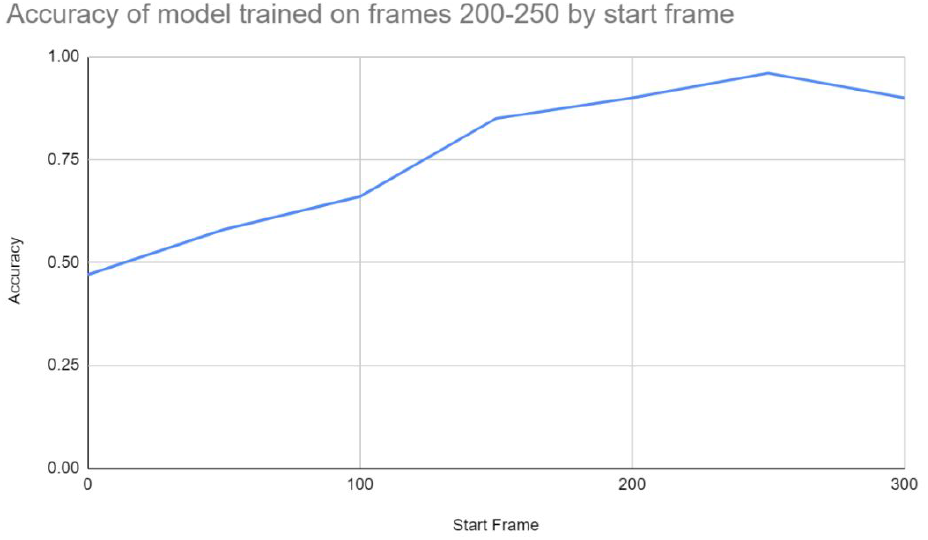
Accuracy of the limited-time model across all held-out time points data. These held-out test accuracy results (y-axis) were aggregated by time (x-axis) into five groups.

### 3.2. Explainable AI techniques reveal differences in T cell interactions with cancer cells under genetic perturbation

To better understand the differences in behavior across genetic perturbations, we applied the Grad-CAM technique^10^ to both full- and restricted-time models and testing with held-out validation data. For an individual sample’s prediction, Grad-CAM combines the gradients of the model’s weight to calculate the influence of each pixel feature to the prediction. These values generate a “feature importance heatmap” that identifies the most important regions of an image for classifications.

We analyzed the output of Grad-CAM across different time points and different conditions to identify changes recognized by CNN classifiers. For the model trained on all time points, we observe that Grad-CAM highlights interactions between cancer cells and T cells, focusing its attention on the cellular aggregates to recognize CUL5KO (Figure 3). In the RASA2KO Grad-CAM visualizations, on the other hand, the highlighted regions are focused almost exclusively on the areas between cells and cellular aggregates (Figure 3). Moreover, we observed that the highlighted regions in the SHKO group seem to be distributed randomLy, but each time focused on individual cancer cells (Figure 3). These Grad-CAM visualizations suggest specific characteristics of behavior of each of the three experiments.

**Fig 3.**
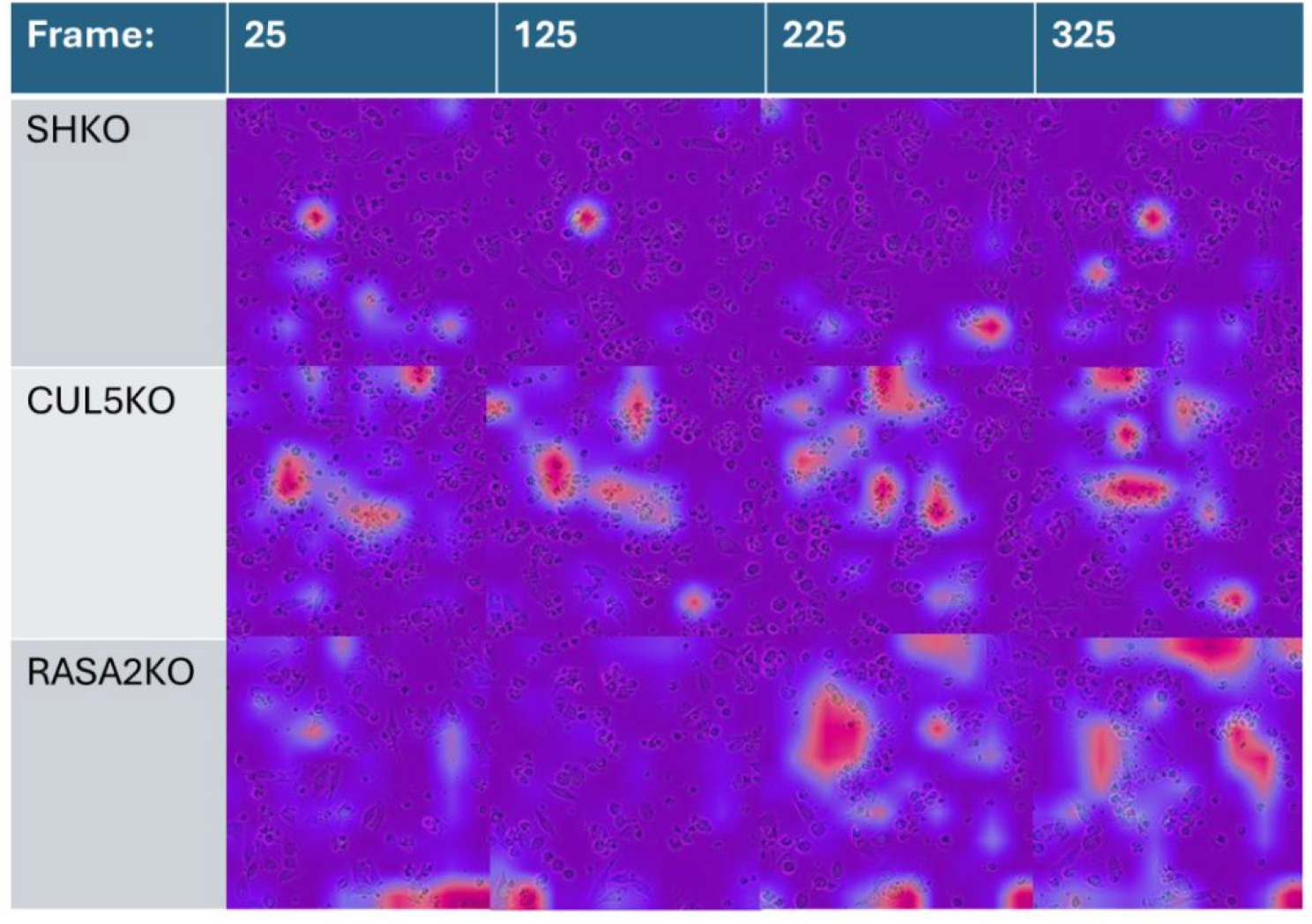
Grad-CAM importance scores for the limited-time model across condition and time on held-out images. Frame label indicates both the true label and associated Grad-CAM class label. The purple areas represent the lowest impact areas, blue represents the medium impact, and red represents the highest impact. This color gradient is consistent for all of the Grad-CAM visualizations throughout this paper.

We quantified the enrichment of these patterns on a small scale in the three experiments by manually annotating the number of healthy and interacting T cells in the highlighted regions of each type on the frame interval 150-160 in the second quadrant from the full time frame model in the held-out images. Across all three sets of heatmaps, the CUL5KO Grad-CAM heatmap highlights the interacting cancer cells at a higher rate than the SHKO and RASA2KO Grad-CAM heatmap (Table 3). This suggests that the difference between CUL5KO and RASA2KO behavior is that CUL5KO T cells accelerate the rate of cancer cell-T cell interactions and the formation of T cell aggregates around a cancer cell.

**Table 3.** Number of healthy and interacting cancer cells from ten images between frames 150 and 160 highlighted by Grad-CAM on the full time frame model.

| Genetic perturbation | Non-interacting cancer cells | Interacting cancer cells |
| --- | --- | --- |
| SHKO | 10 | 5 |
| CUL5KO | 2 | 32 |
| RASA2KO | 12 | 7 |

To better understand differences between the full-time and limited-time models, we compared the Grad-CAM visualizations at the same time from the same position. We use to illustrate one input frame from the CUL5KO held-out data at time point 220,880 minutes post culture (Figure 4). Both sets of Grad-CAM images focus on interacting cancer cells—which often appear as large T cell aggregates that hide the seed cancer cell—but the interacting cancer cells they highlight are often different ones (Figure 4). The 200-249 frame model focuses more on regions of the image without aggregates to inform its decision, indicating that overall T cell/cancer cell coverage is an important signature during this 50-frame time window. Although both visualizations appear to focus on similar proportions of the image, the 200-249 frame model’s heatmap has a larger area of limited attention across the full image (Figure 4).

**Fig 4.**
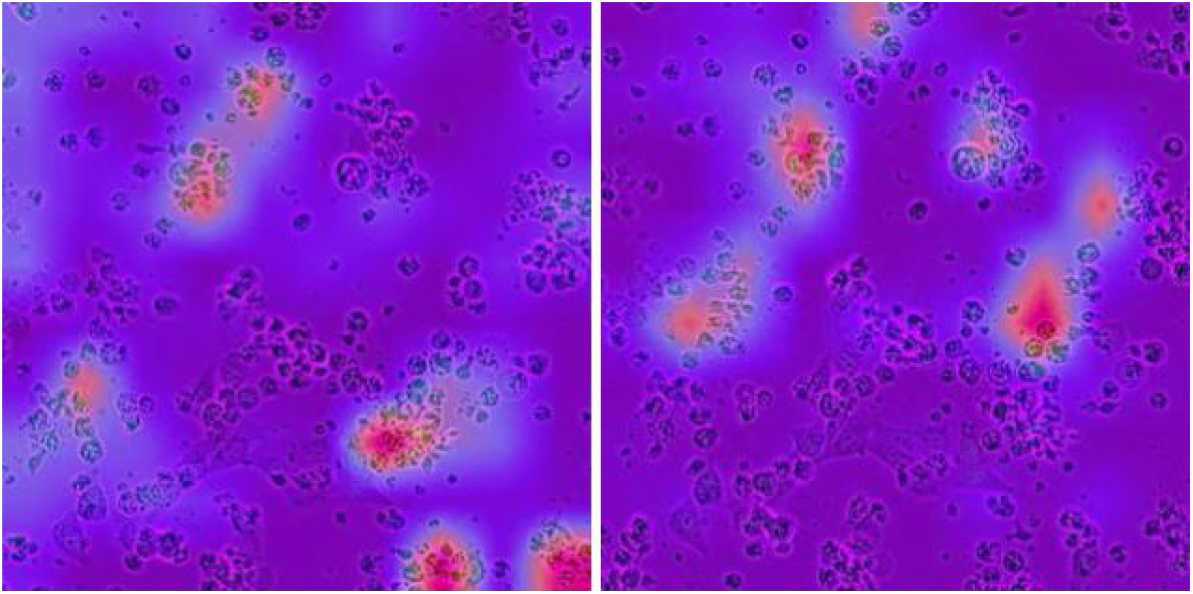
Grad-CAM visualizations of frame 220 of the held out CUL5 images for the 200-249 frame model (left) and full 350 frame model (right).

To more broadly interrogate the influences on the limited-time frame model, we used Grad-CAM to visualization importance heatmaps across the three different genetic perturbation on held-out frames (Figure 5). Like the full-time model (Figure 3) the limited-time model focuses on interacting cancer cells, which we define as T cells adjacent to or overlapping with cancer cells in the CUL5KO held-out frames. The limited-time model, however, has more diffused highlighted regions of importance for predicting all three conditions than the full time model, capturing most of the cells. This suggests that *total cell coverage*, or the proportion of the area of the image covered by cells, is a more defining signature of the CUL5KO limited-time model than the full-time model. When comparing shorter time intervals, CUL5KO can be characterized by its total cell coverage, but, over longer intervals, the specific interactions between cells proves to be the most important distinguishing feature.

**Fig 5.**
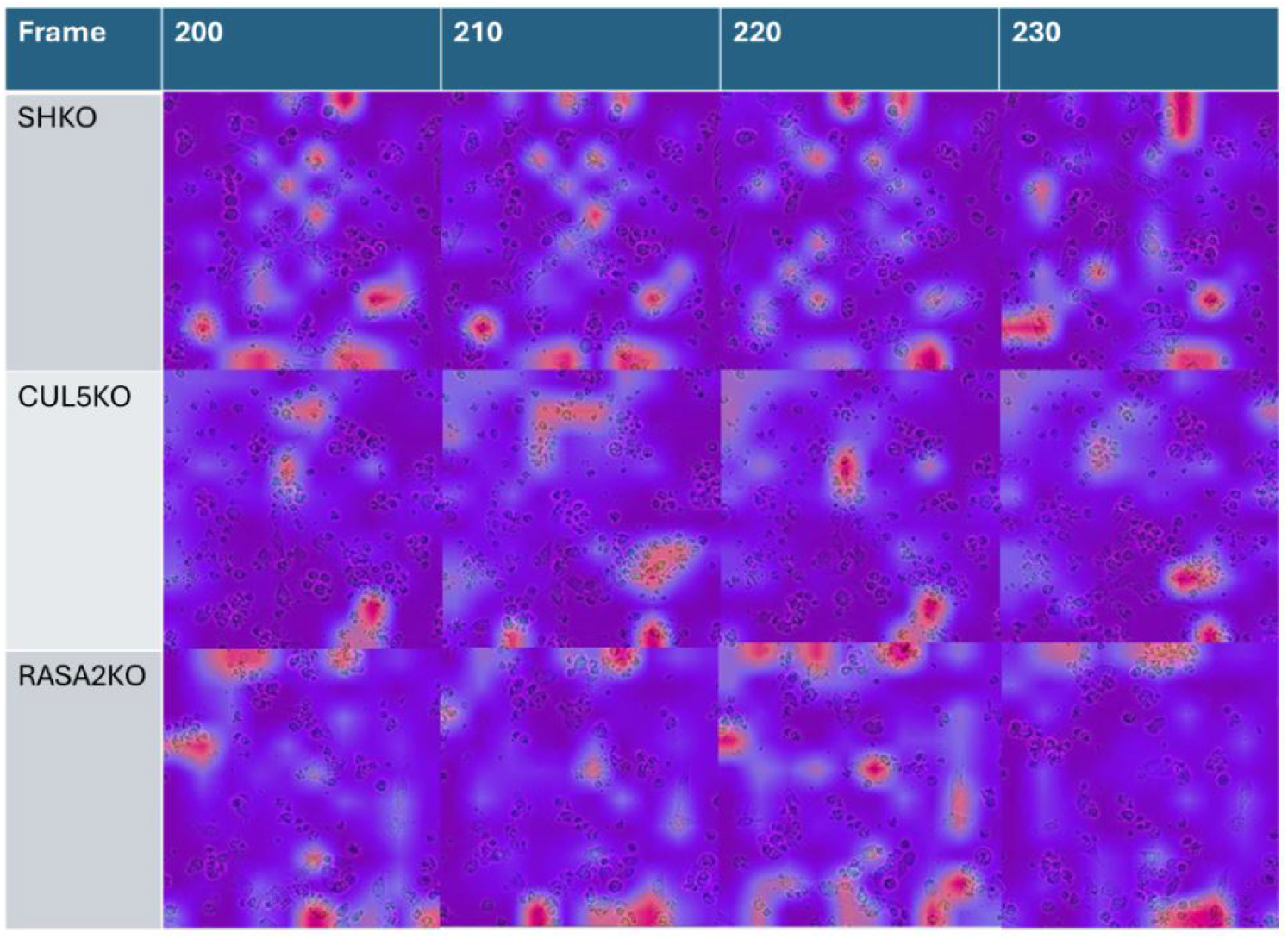
Grad-CAM importance scores for the limited-time model across condition and time on held out images. Frame label indicates both the true label and associated Grad-CAM class label.

Taken together, our findings indicate that the model trained on later limited time frames takes a larger proportion of the image into account when performing classification, whereas the full model focuses on more limited regions of the image. The greater spread of “attention” and the focus on multiple cancer cells and T cells in the limited-time model suggests that the model is effectively counting the number of cells to make a prediction; at limited time points, given the known differences in killing progression, this featurization would be effective for separating the classes as indicated by the limited-time model’s accuracy. In contrast, the model trained on the full data cannot rely on the number of cells to differentiate genetic perturbations, and so focuses more on a small number of cell interaction regions to distinguish the knock-outs.

## 4. Conclusion

Our work analyzes the behaviors of CRISPR-modified TCR T cells interacting with cancer cells in live-cell imaging studies. Most studies count the total area covered by cancer cells across time to characterize the cancer cell killing efficacy of the modified T cells, ignoring the behavior changes in T cell, cancer cells, and their interactions. We identified specific changes in cell behavior across the three experiment types by using Grad-CAM,^10^ a visual explainable AI technique, to understand how a deep learning classifier would differentiate the experimental conditions from live-cell images, highlighting the behavioral changes in modified T cells beyond simple cancer cell death rate.

By using Grad-CAM to analyze classification models trained on three types of modified TCR T cells, we found that the amount of T cell or cancer cell coverage differentiates CUL5KO and RASA2KO modified T cells when comparing similar time points. We showed that cell aggregation behavior is a reliable differentiating characteristic to distinguish CUL5KO experiments from the others, and these CUL5KO experiments tend to have consistent T cell aggregates around cancer cells. We showed that larger empty spaces, suggesting a combination of larger (and fewer) cellular aggregates plus better cancer cell killing, distinguishes RASA2KO experiments from the other two experiments. We found that the safe harbor experiment is defined by no cellular aggregates and substantial coverage of cancer cells, with the T cells both failing to latch on to the cancer cells and furthermore failing to stop their proliferation. We note that coverage plots alone miss these important behavioral signatures.

Our study has a number of limitations, including considering only a single T cell donor and cancer cell line, three genetic modifications, limited replicates, and limited variable titrations of cancer cells to modified T cells. Emerging architectures and pretrained models may improve accuracy relative to the ResNet architecture used here. The importance maps from Grad-CAM are coarse regions over the image, and sometimes the difference between knockouts could be hard to qualitatively observe. Grad-CAM is one of many interpretation approaches, and alternatives such as saliency maps or Shapley Additive Explanations may provide different features of interest. The lack of differences between knockouts may also be an interesting indication of a lack of distinct mechanism changes that would limit downstream efficacy.

However, as a proof of concept, this analysis pipeline for future live-cell imaging experiments will open the door to a more sophisticated interpretation of modified T cell behaviors. We found that existing tools and pretrained image models like ImageNet are effective at classifying biological image samples when fine tuned using live-cell imaging frames. We observed that fine tuning on frames from a wide stretch of time increases the models’ attention on individual cellular dynamics, while fine tuning on short time samples later in the experiment will use more characteristic image features for classification.

Overall, we demonstrated that explainable AI techniques are a practical tool for interrogating and understanding biological dynamics from live-cell image, and we developed a framework for studying these dynamics in general live-cell imaging data. Future work pushes our methods towards the clinic. By characterizing the complex behaviors of these possible T cell modifications, we hope to more rapidly identify T cell therapies for broad ranges of cancers, both liquid and solid. Our interpretable classifiers specifically can be used by decision-making AI methods to prioritize specific T cell therapies for new cancer patients by predicting the response of that individual tumor to each type of therapy, and selecting the most effective therapy.

## Acknowledgements

M.B., A.V., and B.E.E. were funded by NIH/NCI 5U2CCA233195, NIH/NHGRI R01 HG012967, the Parker Institute for Cancer Immunotherapy (PICI), and NIH/NHGRI R01 HG013736. J.C. was supported by NIH/NCI K08, 1K08CA252605-01, a Burroughs Wellcome Fund Career Award for Medical Scientists, the Lydia Preisler Shorenstein Donor Advised Fund, and the Parker Institute for Cancer Immunotherapy, PICI. B.E.E. is a CIFAR Fellow in the Multiscale Human Program. B.E.E. is on the SAB for ArrePath Inc, Crayon Bio, and Freenome; she consults for Neumora.

